# Splicing regulation by RS2Z36 controls ovary patterning and fruit growth in tomato

**DOI:** 10.1101/2025.01.31.635986

**Authors:** Stavros Vraggalas, Remus R E Rosenkranz, Mario Keller, Yolanda Pérez-Pérez, Samia Bachiri, Kerstin J Zehl, Jessica Bold, Stefan Simm, Arindam Ghatak, Wolfram Weckwerth, Leila Afjehi-Sadat, Palak Chaturvedi, Pilar S Testillano, Michaela Müller-McNicoll, Kathi Zarnack, Sotirios Fragkostefanakis

## Abstract

Fruit growth is mediated by cell division and expansion. In tomato, the model for fleshy fruit development, both processes are tightly linked to changes in gene expression, including transcriptional regulation and RNA processing. While several transcription factors are implicated in fruit developmental programs, the role of splicing regulators remains largely unexplored. Expression profiling of splicing-related genes revealed expression patterns. The serine/arginine-rich splicing factor RS2Z36 is expressed in ovaries and during early fruit development. Loss-of-function mutations in RS2Z36 result in ovaries with altered patterning and in smaller, ellipsoid fruits. *rs2z36* mutants display elongated pericarp cells along the longitudinal axis of pre-anthesis ovaries, indicating that RS2Z36-dependent expansion patterns are established before anthesis. RNA-seq uncovered widespread alternative splicing of genes across diverse biological processes, while proteome analysis revealed altered protein abundance and peptides derived from novel splice variants. In addition, *rs2z36-1* pericarps show increased deposition of LM6-recognized arabinan and AGP epitopes. Together, these findings identify RS2Z36 as a regulator of ovary and fruit development and highlight a previously underappreciated role for splicing control in shaping early fruit morphology.

## Introduction

The growth of fleshy fruits depends on tightly coordinated cell division and expansion programs. Tomato (*Solanum lycopersicum*) is a model for studying fleshy fruit development. Fruit morphology is genetically determined, and core loci that influence fruit size and shape, contributing to the diversity seen in modern tomatoes through domestication and breeding, have been identified (Paran and Van Der Knaap, 2007). *LC* and *FAS* loci in tomato, encoding orthologs of *Arabidopsis WUSCHEL* (WUS) and *CLAVATA3* (CLV3), regulate locule number by altering the spatial and temporal expression of *SlWUS*, leading to multilocular and enlarged fruits compared to wild progenitors (Cong et al., 2008; Muños et al., 2011; Chu et al., 2019). Mutations in *LC* and *FAS* expand *SlWUS* expression in floral buds, increasing meristem size and positively influencing fruit growth through transcriptional regulation of genes involved in meristem maintenance and floral organ development (Chu et al., 2019). The genes *SUN* and *OVATE* control elongated fruit shape (Xiao et al., 2008). *SUN* codes for an IQ67 protein involved in calcium signalling and trafficking through interactions with calmodulins (CaMs) and kinesin-light chain proteins (Abel et al., 2013; Clevenger et al., 2015). Its mutation, caused by Rider retrotransposon-mediated duplication, leads to high *SUN* expression and enhanced longitudinal growth (Xiao et al., 2008; Jiang et al., 2009). *OVATE* encodes a transcriptional repressor; a mutation introducing a premature termination codon (PTC) produces elongated fruits (Liu et al., 2002; Wang et al., 2007). OVATE is also linked to microtubule organization and cell division through interactions with TONNEAU1 Recruiting Motif proteins (Snouffer et al., 2020).

Tomato fruit growth, from ovary development to the mature green stage, involves overlapping programs of cell division and expansion (Renaudin et al., 2017; Penchovsky and Kaloudas, 2023). Pericarp cell division occurs mainly within 8–10 days post-anthesis, though it can extend to 25 days. Expansion of mesocarp cells then drives dramatic fruit enlargement, with expansion orientation affecting pericarp growth and thickness (Musseau et al., 2017). Enlargement is driven by vacuolar expansion, cytoplasmic growth, and endoreduplication (Cheniclet et al., 2005). *SlCCS52A* facilitates the transition from mitotic to endocycles, allowing higher ploidy and increased cell size (Mathieu-Rivet et al., 2010). The cytoskeleton provides structural support for vesicle trafficking and cell wall remodelling (Mauxion et al., 2021), while *SUN* and *OVATE* influence cytoskeletal organization and cell division patterns, affecting fruit shape (Wang et al., 2019).

Hormonal signalling plays a central role in fruit development (Azzi et al., 2015). Auxins regulate fruit set and early growth, with *SlIAA17* modulating endoreduplication-linked expansion (Su et al., 2014). Gibberellins, controlled by *SlGA2ox1*, stimulate growth during early stages (Chen et al., 2016). Metabolic processes underpin these phases, supplying energy and precursors. Sugar metabolism, regulated by *SlLIN5* and *SlHXK1*, ensures carbohydrate availability for wall biosynthesis and energy production (Zanor et al., 2009; Li et al., 2023).

The ovary-to-fruit transition involves major transcriptome changes, including 700–2800 differentially expressed genes in pericarp tissue during the first 5 days post-anthesis (Pattison et al., 2015; Zhang et al., 2016). Hundreds of transcription factors show differential expression across stages. In contrast, alternative splicing (AS) has been less studied, although ∼65% of tomato protein-coding genes undergo AS (Clark et al., 2019). AS has been linked to ripening and quality attributes (Chen et al., 2019; Colanero et al., 2020; Zhou et al., 2022), but its role in early development is poorly defined. RNA-seq analyses reveal numerous stage-specific AS events, including in auxin response factor (ARF) genes mediating auxin signalling (Zouine et al., 2014; Sun and Xiao, 2015; Wang et al., 2016).

Pre-mRNA splicing is a crucial step in mRNA maturation, in which introns are removed and exons are joined within a pre-mRNA transcript [citation]. This process is mediated by the spliceosome, a large RNA-protein complex composed of U1, U2, U4/U6, and U5 snRNPs (Reddy et al., 2013; Meyer et al., 2015). Each snRNP contains an snRNA bound by Sm-like proteins, providing structural stability and assembly function (Perea-Resa et al., 2013). AS describes the selective usage of splice sites, generating multiple mRNA variants from a single gene. Each snRNP contains an snRNA bound by Sm-like proteins, providing structural stability and assembly function (Perea-Resa et al., 2013). Alternative splice site selection can enhance proteome diversity through the generation of isoforms, and can regulate mRNA levels, e.g. through the formation of premature stop codons (PTCs) and subsequent nonsense-mediated decay (NMD) (Reddy et al., 2013).The main AS types in plants are intron retention, exon skipping, and alternative 3′ and 5′ splice sites (Staiger and Brown, 2013).

Splice site choice is influenced by serine/arginine-rich (SR) proteins and heterogeneous nuclear ribonucleoproteins (hnRNPs) by binding to cis-elements and recruiting spliceosome components. SR proteins are characterized by N-terminal RNA recognition motifs (RRMs) and a C-terminal arginine/serine-rich (RS) domain (Barta et al., 2010; Howard and Sanford, 2015; Wegener and Müller-McNicoll, 2019). Studies in *Arabidopsis* show that SR proteins regulate development, reproductive and vegetative organ growth, and flowering (Lopato et al., 1999; Ali et al., 2007; Zhang and Mount, 2009; Stankovic et al., 2016; Yan et al., 2017). In tomato, RS2Z35 and RS2Z36 regulate heat stress–sensitive AS and contribute to thermotolerance (Rosenkranz et al., 2024). Members of the SR gene family also show distinct expression patterns during development (Rosenkranz et al., 2021), but their roles in fruit development remain unexplored.

Here, we performed global expression profiling of splicing-related genes in tomato fruit tissues and stages, revealing tissue- and stage-specific expression. RS2Z36, highly expressed in ovaries and early fruit, was studied in CRISPR/Cas9 mutants. Both lines produced smaller, more ellipsoid fruits compared to wild type. Integration of transcriptome and proteome data uncovered RS2Z36-dependent splicing changes potentially linked to fruit growth and morphology. We propose that RS2Z36 regulates AS during ovary development, fine-tuning transcript and protein diversity required for fruit size and shape determination. Our study underscores RNA splicing as a key mechanism shaping developmental traits in tomato.

## Results

### Expression of SR proteins during tomato fruit growth and development

Previous studies have shown that many multi-exonic genes expressed during tomato fruit growth and development undergo AS (Sun and Xiao, 2015; Wang *et al*., 2016). To identify molecular mechanisms that regulate RNA splicing during this developmental process, we examined the expression profile of genes coding for splicing regulators using the Tomato Expression Atlas (TEA) database (https://tea.solgenomics.net/) which provides information on different tissues of fruits from anthesis to ripe stage (Supplemental Figure 1; Shinozaki *et al*., 2018). The set of 198 genes included orthologues of *Arabidopsis* splicing-related genes including spliceosomal components such as snRNPs as well as associated factors such as hnRNPs and SR proteins (Supplemental Table 2; Rosenkranz *et al*., 2021). The samples included various fruit tissues, and for each, a series of developmental stages as shown in the right panel of Fig. 1A. A co-expression analysis resulted in six clusters, each comprised of 21 to 57 genes (Fig. 1A-B). In cluster 1, 38 genes are expressed at higher levels at anthesis and then gradually decrease during fruit development, while those of cluster 5 exhibit a strong transient increase with 5 days from anthesis. Twenty-four genes of cluster 2 show a gradual increase from anthesis which peaks at mature green (MG) fruits and then drop during fruit ripening, while 57 genes in cluster 6 show a similar induction but are maintained at high expression levels throughout fruit ripening (Fig. 1A-B). Twenty-one and 25 genes in cluster 3 and 4, respectively, exhibit a steady expression, while those of cluster 4 show a strong increase specifically in seed embryos (Fig. 1A-B).

**Figure 1.**
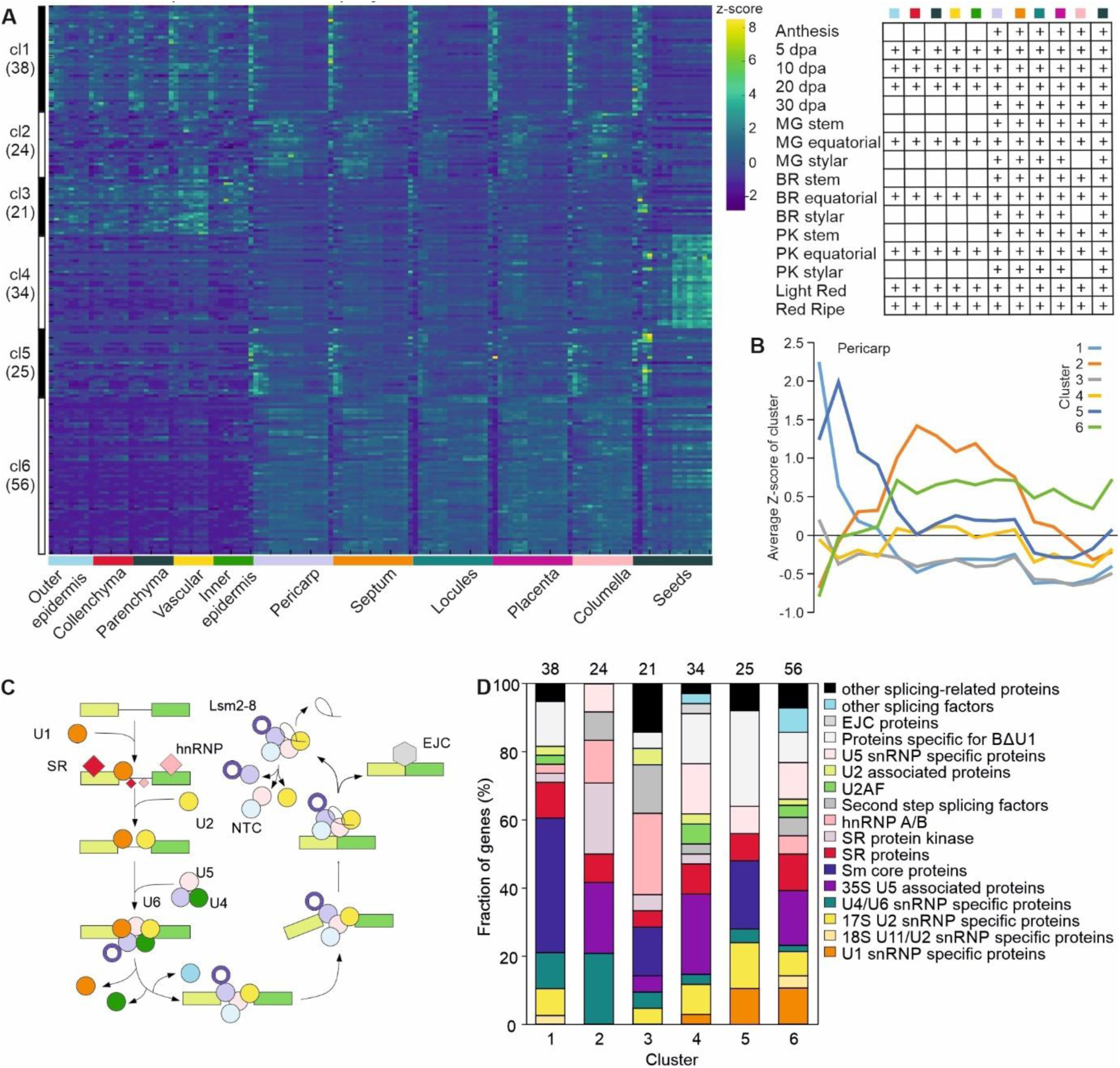
Expression analysis of tomato RNA splicing related genes in tomato tissues during development and ripening. (A) Heat map and clusters of RNA splicing related genes in different tomato tissues and developmental stages. Data have been acquired from the Tomato Expression Atlas database (Shinozaki *et al*., 2018). The panel on the right indicates the different stages of each tissue that were included in the analysis. dpa: days past anthesis. (B) Mean expression of genes from each cluster in pericarp tissue during fruit development. The x-axis refers to the samples shown in the panel above for pericarp. (C) Overview of pre-mRNA splicing regulation. The scheme is adopted from (Meyer *et al*., 2015). (D) Composition of gene clusters.

The stage-specific expression patterns of genes encoding RNA splicing factors and associated proteins suggest that distinct splicing complexes might be selectively assembled to regulate splicing in a developmental stage-specific manner (Fig. 1C). For instance, U1 snRNP-coding genes are predominantly found in clusters 4, 5, and 6, whereas U4/U6 snRNP-coding genes are mainly grouped in clusters 1 and 2 (Fig. 1D). Among the gene families encoding core AS regulators, SR genes exhibit broad distribution across all clusters, indicating their involvement throughout various stages (Fig. 1D). In contrast, hnRNP genes are largely concentrated in clusters 2 and 3, suggesting a more restricted stage-specific role (Fig. 1D). These patterns highlight the dynamic and coordinated regulation of splicing machinery components during different stages of development.

### RS2Z36 is involved in fruit growth

We focused on cluster 5, as it contains genes showing a peak in expression in the narrow period of 5-10 dpa (Supplemental Figure 2A), coinciding with the end of cell division and the beginning of cell expansion (Azzi *et al*., 2015). Among the genes of cluster 5, Solyc09g005980 codes for RS2Z36, a protein that belongs to the plant specific clade of SR proteins and was previously shown to be important for thermotolerance by regulating heat stress sensitive AS in leaves and seedlings, while a mutant line of this gene does not have any phenotype in vegetative tissues under control conditions (Rosenkranz *et al*., 2021, 2024). Interestingly, RS2Z35, the paralogue of RS2Z36, belongs to cluster 6, thereby having a different expression profile, suggesting that the two genes might have distinct roles in the regulation of AS during fruit development.

To examine whether RS2Z36 plays a role in tomato growth and development, we utilized CRISPR/Cas9 mutants with two guide RNAs (gRNAs) targeting the RRM coding region in exons 2 and 4 (Fig. 2A). The *rs2z36.1* line has an insertion of one nucleotide, generating a premature termination codon (PTC) in the RRM coding region (Fig. 2A). Both mutant lines carry a second deletion of one nucleotide in the 3’-half of the RRM coding region, resulting in a PTC in *rs2z36.2* as well (Fig. 2A). Expression analysis by qRT-PCR revealed that the transcript levels of *RS2Z36* were reduced in the mutant lines compared to the WT (Fig. 2B). This indicates that the generation of a PTC in both mutants likely results in the degradation of the *RS2Z36* RNA by NMD.

**Figure 2.**
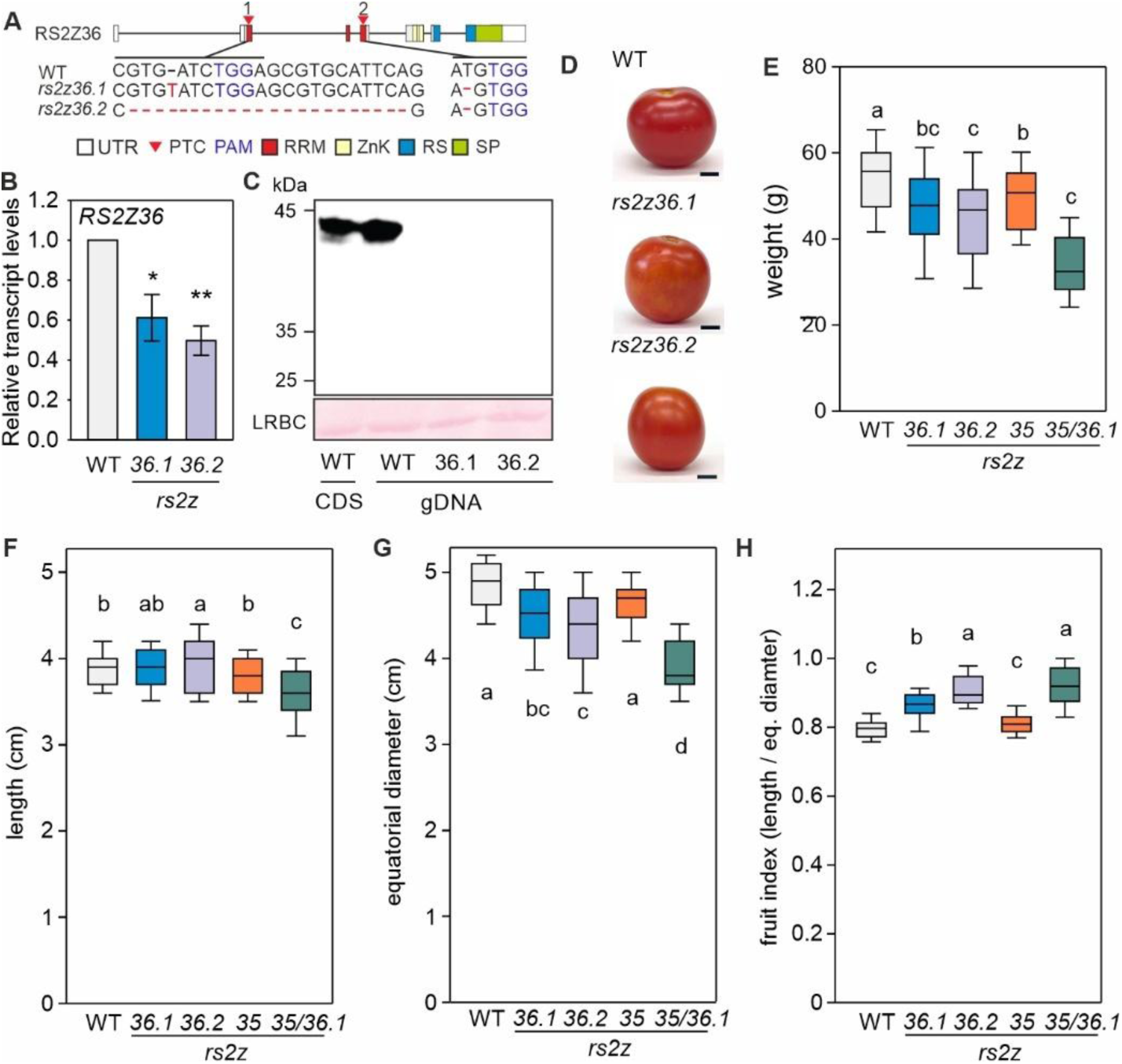
Mutations of the *RS2Z36* gene affect fruit size and shape. (A) CRISPR/Cas9-mediated mutations in two independent tomato lines. Boxes indicate exons, and colours coding regions for protein domains. UTR: untranslated region, PTC: premature termination codon, RRM: RNA-recognition motif, ZnK: zinc knuckle, RS: serine/arginine-rich region, SP: serine/proline-rich region. Arrows indicate the gRNA targeting region and the blue nucleotides the PAM sequence. (B) Expression of *RS2Z36* transcript levels in leaves of WT and *rs2z36* mutants based on qRT-PCR. Bars indicate the mean of three biological replicates ±SEM. Asterisks indicate statistically significant differences based on a t-test (* *p* < 0.05, ** *p* < 0.01). (C) Immunodetection of WT and mutated RS2Z36 in tomato protoplasts. Protoplasts were transformed with DNA carrying the coding sequence of the WT RS2Z36 (CDS), or the gDNA from WT or the *rs2z36* mutants. All constructs were N-terminally tagged with a 3xHA peptide. Proteins were detected with an αHA antibody. The lower blot indicates the large subunit of RuBisCO after Ponceau staining. (D) Representative images of WT and *rs2z36* mutant fruits at ripening stage. Bar = 1 cm. (E) Weight, (F) length, (G) equatorial dimeter, and (H) fruit index of ripe WT, *rs2z36.1*, *rs2z36.2*, *rs2z35* and the double *rs2z35 rs2z36.1*. Box plots show the median from 15-20 fruits. Statistically significant differences are based on ANOVA tests with post-hoc Duncan’s multiple range test (*p* < 0.05).

To examine whether RS2Z36 protein could be synthesized in the *rs2z36* mutant lines, as an antibody against RS2Z36 is not available, we used in-frame tagged RS2Z36-expressing constructs. For this *in vitro* assay, the genomic region of the RS2Z36 gene from the start to the stop codon from WT and both mutant lines were cloned under the control of CaMV35S promoter and N-terminally fused in frame with three consecutive human influenza hemagglutinin (HA) peptides. Following plasmid transfection of tomato protoplasts, immunoblot analysis with an anti-HA antibody showed that only the WT gRS2Z36 protein could be detected, further supporting the assumption that the mRNAs of both mutants are targeted for NMD and are not translated into protein (Fig. 2C).

The mutant plants were phenotypically similar to the WT plants at the vegetative stage under control conditions but showed noticeable differences in fruit morphology (Fig. 2D). The ripe fruits of *rs2z36.1* and *rs2z36.2* mutants had 14.2 and 16.1% lower weight compared to WT, respectively (Fig. 2E). Both lines have similar fruit length to WT but a reduced equatorial diameter by 6.7 and 10.2% compared to WT in *rs2z36.1* and *rs2z36.2, respectively* (Fig. 2F-G). This resulted in fruits of more elliptical shape (Fig. 2H) compared to WT, as indicated by the 8.8-12.2% higher index of the mutant fruits compared to WT (Fig. 2H). To investigate whether this phenotype is specific for RS2Z36, we examined a CRISPR knockout mutant line for RS2Z35 as well as a double knockout line (*rs2z25 rs2z36.1*) which have been previously described (Rosenkranz *et al*., 2024). While the fruits of *rs2z35* plants did not appear to have an altered shape (Fig. 2E-H), the fruits of the double mutant were reduced in their length and diameter and showed a fruit index similar to those of the single *rs2z36* mutants (Fig. 2E-H). We did not observe any significant effect on the timing of the ripening process (not shown). Overall, these results suggest an involvement of RS2Z36 in fruit morphology.

Knowing that RS2Z36 is mainly expressed during the first days after anthesis (Fig. 1B), we performed qRT-PCR to examine its expression in ovaries collected from flowers 2 days before anthesis and at anthesis, the latter as a common reference sample to the TEA dataset (Figure 1A). RS2Z36 expression is approximately 40% higher in ovaries before anthesis than at anthesis (Supplemental Figure 2B). Consequently, we examined the size and shape index of the ovaries from flowers at various stages 4 days before to 4 days after anthesis. At all time points, the ovaries of the *rs2z36* lines had a more elliptical shape compared to WT (Fig. 3A-B). To examine whether this phenotype is associated with alterations in cellular morphology, we performed longitudinal sections and examined the shape index of mesocarp cells from ovaries of flowers at anthesis (Fig. 3C-D). For this, measurements were performed on cells from the equatorial region of the mesocarp, and the maximum length of the equatorial diameter was considered. In both mutant lines, the mesocarp cells were more elongated compared to the cells in the WT ovaries, suggesting that the elliptical ovary and fruit shapes are a consequence of cell elongation caused by *RS2Z36* mutations (Fig. 3C-D).

**Figure 3.**
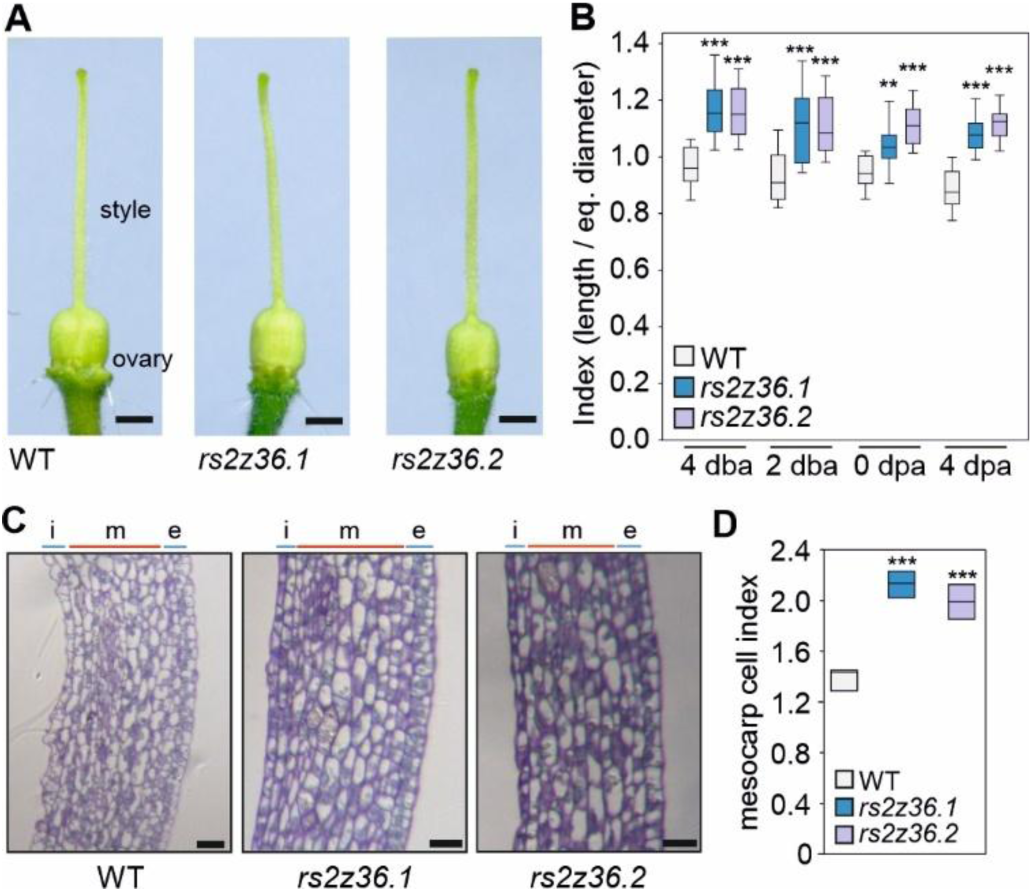
The shape of ovaries from *rs2z36* mutant lines is more elliptical compared to wild type. (A) Representative flowers from WT and *rs2z36* lines. Bar = 2 mm. (B) Ovary index at 4 and 2 days before anthesis (dba), at anthesis (0) and at 4 days post anthesis (dpa). Box plots are based on 15-20 fruits for each sample. (C) Representative images of the equatorial region from toluidine blue stained longitudinal sections of fruits from WT and *rs2z36* mutants at anthesis. Bar = 25 μm. (D) Cell index from the mesocarp (m) region (cell layer between 3 cell layers of epicarp (e) and inner cells of the endocarp (i), from 4-6 independent fruits each (>40 cells from each fruit from 3 sections). Statistically significant differences between the WT and the mutants are indicated by two (*p* < 0.01) or three asterisks (*p* < 0.001) based on T-test.

### RS2Z36-dependent alternative splicing in tomato ovaries

As the phenotype of the altered fruit morphology is already present in ovaries before anthesis, at a stage where RS2Z36 is highly expressed, we performed a transcriptome analysis on RNA isolated from ovaries at 2 days before anthesis (dba) from wild type and the *rs2z36.1* line. We identified 20,621 genes with TPM ≥ 0.5 in at least one line (WT or *rs2z36-1*), with the cutoff consistently met across all replicates. MAJIQ was used to analyse AS events (delta percent selected index [|ΔPSI|] ≥ 0.05 and probability changing [*P*_(|ΔPSI| ≥ 0.02)_] ≥ 0.9), as previously described (Rosenkranz *et al*., 2024; Supplemental Table 3). Only binary AS events were analysed, namely IR, CE, A5SS and A3SS, while complex events were excluded. In total, we found 240 differential AS (DAS) events occurring in 230 genes (Fig. 4A). Most of the events belong to alternative A3SS (93 DAS events), followed by IR, A5SS and CE with 56, 50 and 42 events, respectively (Fig. 4A).

**Figure 4.**
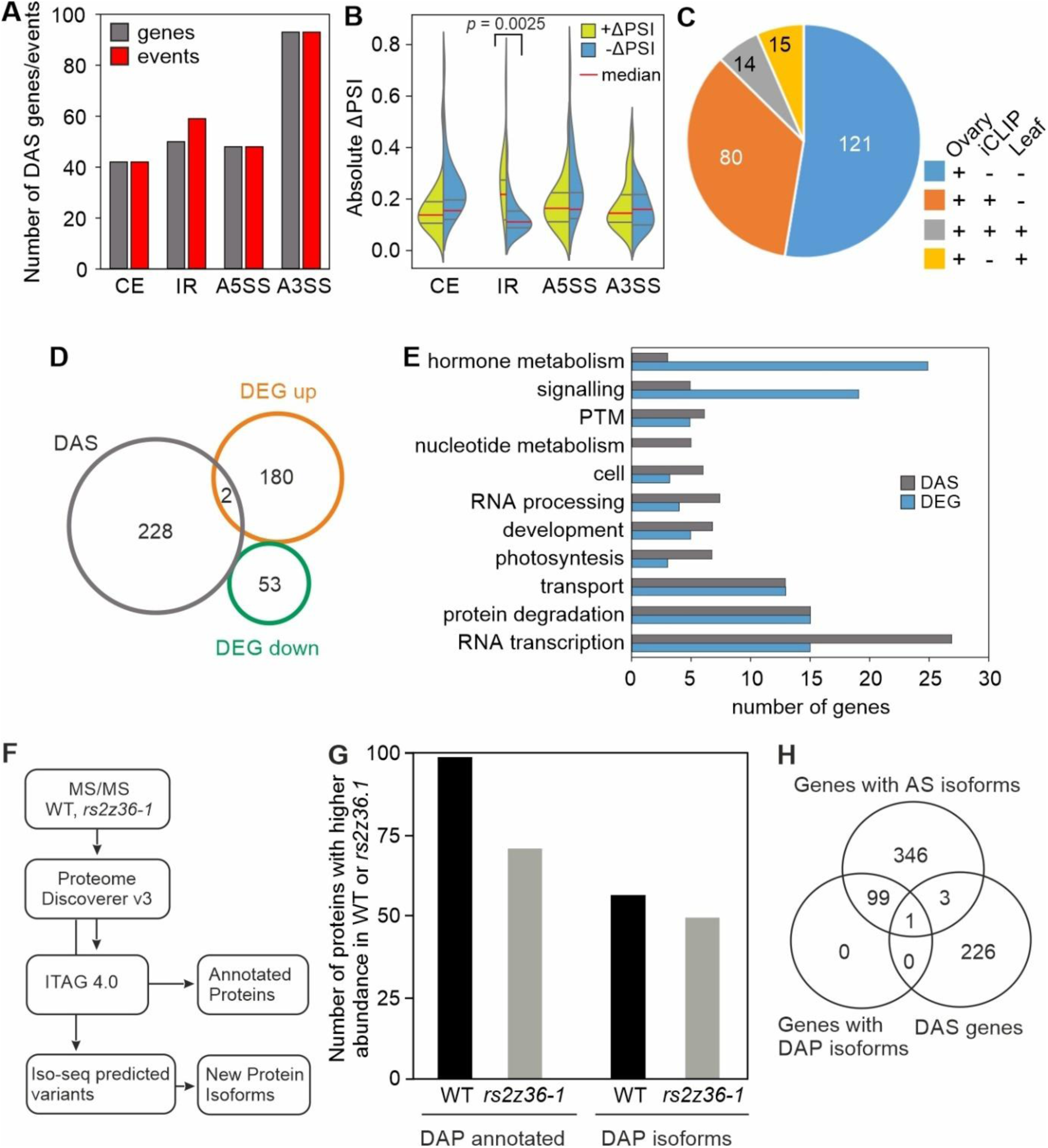
Regulation of AS and gene expression by RS2Z36 in tomato ovaries. (A) Number of DAS events and genes in four main AS categories. CE: cassette exon, IR: intron retention, A5SS and A3SS: alternative 5’ and 3’ splice site selection, respectively. (B) Violin plot of the absolute ΔPSI values of DAS events. Red line is the median and the grey lines above and below are the quartiles. (C) Pie chart showing the overlap of DAS genes identified in ovaries with DAS genes and iCLIP targets reported in leaves by Rosenkranz *et al*. (2024). (D) Number of up- and downregulated DEGs and comparison to DAS genes. (E) List of top biological processes that DAS genes and DEGs are involved in. A full list is provided in Supplemental Table 5. (F) Proteome analysis workflow. (G) Number of differentially abundant annotated proteins (DAP annotated) and peptides resulting from AS (DAP isoforms), expressed at higher levels either in the WT or *rs2z36.1* ovaries. (H) Venn diagram of DAS genes and genes corresponding to differentially abundant annotated proteins and isoforms.

For CE, A5SS and A3SS, the ΔPSI values were equally distributed in both directions, indicating that the mutation of RS2Z36 results both in enhancement and suppression of AS in a gene specific context (Fig. 4B). However, for IR, there was a significant shift towards intron splicing, suggesting that for these events, RS2Z36 functions primarily as intron splicing silencer (Fig. 4B).

A comparison of the RS2Z36-dependent DAS genes in ovaries with the RS2Z36-dependent DAS genes in leaves from plants grown under control conditions (Rosenkranz *et al*., 2024) revealed only 29 common DAS genes in both organs, suggesting that RS2Z36 has organ and tissue specific roles in tomato plant development (Fig. 4C). In addition, the iCLIP dataset generated on heat stressed leaves (Rosenkranz *et al*., 2024) was utilized to investigate whether the RNAs of ovary DAS genes are likely to be directly bound by RS2Z36. 41% of the RS2Z36-dependent DAS genes in ovaries are listed as iCLIP targets, suggesting a possible direct regulation of these AS events by RS2Z36 (Fig. 4C).

Analysis of changes in gene expression revealed a total of 235 differentially expressed genes (DEGs), of which 182 were upregulated and 53 downregulated in the *rs2z36.1* compared to WT ovaries (|log_2_FC| >1, padj ≤ 0.05) (Fig. 4D; Supplemental Table 4). Only two genes were detected both as DAS and DEG, namely Solyc07g017530 annotated as “conserved oligomeric Golgi complex subunit 3” and Solyc06g050710 coding for a putative “heavy metal transport/detoxification superfamily protein”.

Gene ontology analysis of DAS genes or DEGs did not yield an enrichment of a particular biological process for the regulated genes (not shown). Based on MapMan annotation, several genes coding for proteins related to RNA transcription, signalling, transport, and protein degradation and modification were among DAS genes and DEGs (Fig. 4E; Supplemental Table 5). We did not find any known tomato fruit growth related genes to be affected by RS2Z36 mutations. However, we identified DAS genes or DEGs coding for ethylene, gibberellin and auxin related proteins, cell wall proteins as well as cyclins, which could potentially be associated with fruit growth (Fig. 4E, Supplemental Tables 3-5).

### Proteome analysis for the identification of protein isoforms

To identify whether differential expression or altered splicing in the mutants results in changes in the levels of specific proteins, we performed a proteome analysis of protein extracts from ovaries of WT and *rs2z36.1* plants. The identified peptides were filtered to retain only those matching annotated proteins in the tomato genome, the products of canonical/constitutive splicing (Fig. 4F). We further performed differential abundance analysis in the *rs2z36.1* ovaries relative to the WT. From 1256 annotated proteins, we identified 165 proteins with significantly altered levels (p ≤ 0.05; differentially abundant proteins; DAP) between the WT and *rs2z36.1* ovaries, with 96 being more abundant in WT and 69 in the mutant (Fig. 4G; Supplemental Table 6). The proteins that were more abundant in WT ovaries included, among others, several ribosomal proteins, four cell wall related proteins (endo-1,4-beta-xylanase, expansin precursor 5, expansin and pectinesterase), as well as SCL29 (Solyc01g005820), a member of the SR protein family (Supplemental Table 6). Two profilin and six histone proteins, among others, were more abundant in *rs2z36.1* ovaries compared to WT.

Furthermore, the peptides that could not be mapped to the annotated tomato proteome were screened against a long-read library of putative tomato protein isoforms derived from novel RNA splice variants based on an in-house Iso-Seq database (ThermoTom unpublished) and other database sources from Sol Genomics Network (SGN) (Hosmani et al. 2019). The RNA splice variant library was generated from RNA of tomato roots, leaves, flower buds, immature and mature green fruits, exposed to 20, 35, 40, 45 °C for one hour (unpublished). Using the long-read library to screen for possible open reading frames (ORFs) that are not corresponding to the annotated proteome allowed us to identify novel splice variants with high accuracy, capturing full-length transcripts and providing a more comprehensive view of the splicing landscape. Considering that it is currently not known which RNA variants are translated, all possible ORFs that are generated by these RNAs were considered. This analysis yielded peptides that are putatively encoded by RNA splice variants of 449 genes found in our library (Fig. 4H, Supplemental Table 7), with 54 peptides being more abundant in WT and 50 in the *rs2z36.1* ovaries, which are further referred to as DAP isoforms (Fig. 4G; Supplemental Table 7). Except for of two peptides that correspond to two splice variants of a single gene, all peptides are predicted to derive from different genes.

To our surprise, only 1 of the peptides identified as DAP isoform derived from an RS2Z36-dependent DAS gene. Solyc12g044860 codes for an ATP-dependent RNA helicase harboring a DEXDc and a HELICc domain (Fig. 5D). Using a combination of short and long read mapping, we were able to assign two additional protein isoforms. A5SS in intron 4 and retention of introns 6-8 result in a shorter protein isoform that carries only the HELICc domain. This variant is predicted to derive from a short ORF in exon 8 and the flanking introns, meaning that this mRNA has very long 5’ and 3’ untranslated regions (UTRs; Supplemental Figure 5). Additionally, we were able to map another peptide to a splice variant identified by the Iso-seq library but not the short-read DAS analysis, which spans exon 5 to exon 7 as a result of a complex splicing event and in this case results in an RNA with a premature termination codon and a long 3’-UTR, which yields a truncated helicase missing the HELICc domain (Supplemental Figure 5).

**Figure 5.**
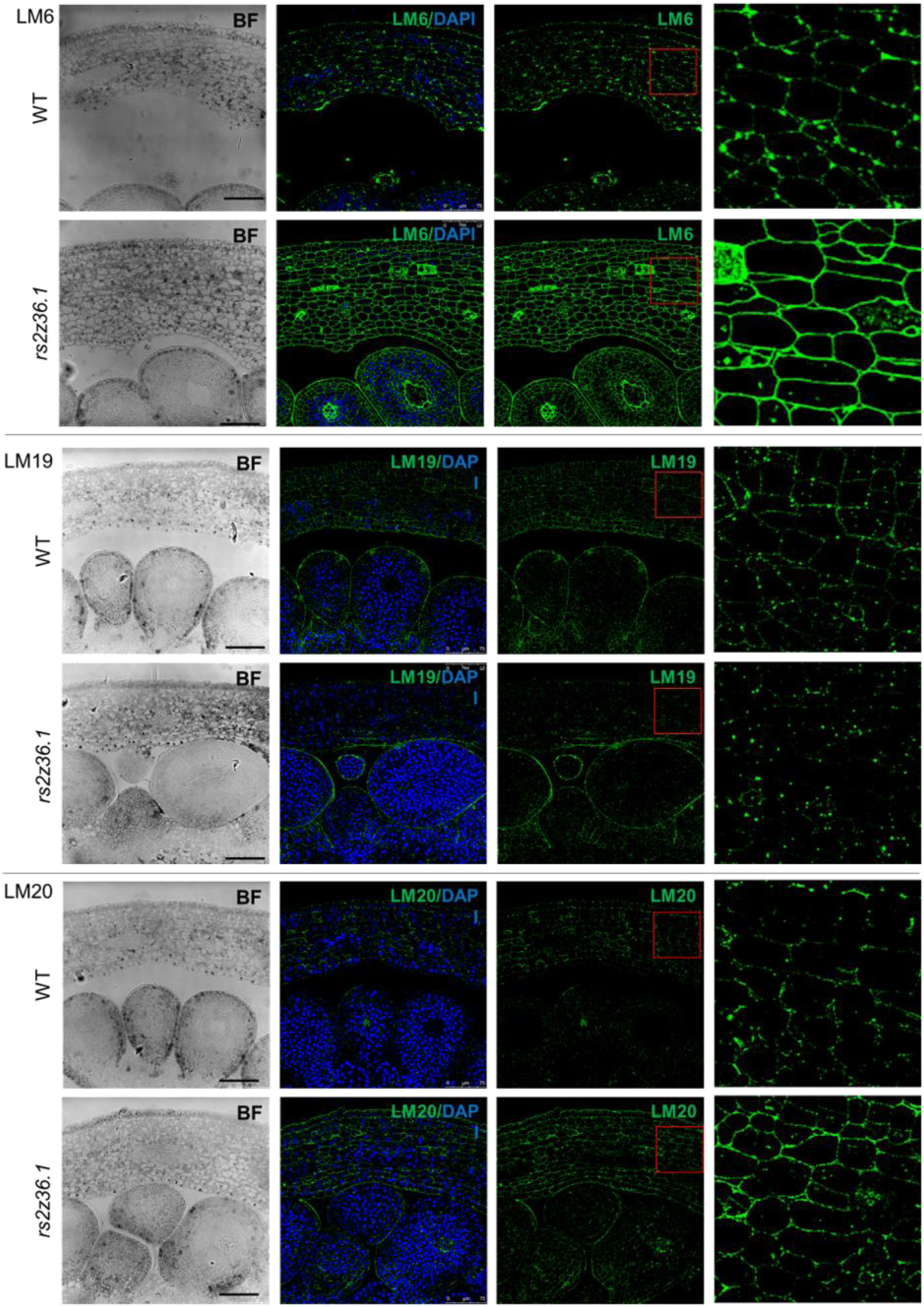
Immunolocalization of AGP epitopes in ovaries at anthesis. Longitudinal sections of ovaries from WT and *rs2z36.1* mutant lines were immunolabeled with LM6 (top) recognizing (1→5)-α-L-arabinan epitopes in rhamnogalacturonan-I (RG-I) and arabinogalactan protein (AGPs), and LM19 (middle) and LM20 (bottom) detecting de-methylesterified and highly methylesterified homogalacturonan (HG) epitopes, respectively. In blue, DAPI staining indicates nuclei. On the left, micrographs of bright field controls are shown. Bars are 75 μm. In the left panel, a magnification of the area indicated by the red box is shown. A negative control is provided in Supplemental Figure 1.

The absence of an overlap of DAS and DAP isoforms, which could also be due to the absence of complex splicing events in our DAS inventory, did not allow us to directly connect changes in AS to specific protein abundance; however, changes in proteome diversity and abundance indicate a potential indirect link. While Gene Ontology analysis did not yield any enrichment of a particular biological process for either the DAP proteins and DAP isoforms, in both lists we were able to find proteins that might be associated with pathways regulating fruit growth and morphology such as cell wall proteins and hormone metabolism (Supplemental Table 8).

### Immunostaining of cell wall components

The transcriptome and proteome analyses revealed several genes as DAS, DEG or DAP that might be associated with the phenotype of elongated pericarp cells, and several of these genes were related to the cell wall. The remodeling of cell wall polysaccharides is a central determinant of tomato fruit growth, driving both cell expansion and tissue architecture. Immunolabeling with well-established monoclonal antibodies provides insights into the distribution and modification of pectic domains during development. LM6 recognizes arabinan side chains of rhamnogalacturonan I (RG-I), which contribute to wall flexibility and hydration and are typically abundant in expanding tissues (Willats et al., 1998). LM19 and LM20 target homogalacturonan (HG) with contrasting methylesterification states: LM19 binds to de-methylesterified HG, associated with calcium-mediated cross-linking and wall stiffening, whereas LM20 detects highly methylesterified HG, linked to wall plasticity and extensibility (Verhertbruggen et al., 2009). Together, these epitopes serve as markers for the dynamic balance between wall loosening and strengthening, reflecting the developmental transitions from early fruit expansion to later stages of wall consolidation and ripening.

We used these three antibodies to detect potential changes in cell wall composition in WT and *rs2z36.1* ovary sections using immunofluorescence. LM6, which recognizes (1→5)-α-L-arabinan epitopes in the arabinan components of rhamnogalacturonan-I (RG-I) and arabinogalactan protein (AGPs), showed a higher immunofluorescence intensity in the mutant ovary sections compared to WT, as well as wider distribution of the signal that homogeneously covered the cell walls in the mutant, while the WT showed a patchy pattern along the cell wall structure (Fig. 5). The increased signal and modified pattern of LM6 labelling suggests enhanced arabinan deposition or changes in its structure in the mutant. Labelling with LM19 and LM20, which detect de-methylesterified and highly methylesterified homogalacturonan (HG), respectively, showed reduced LM19 and increased LM20 labelling, suggesting a shift towards methylesterified HGs (Fig. 5).

## Discussion

Despite the well-established importance of AS in gene regulation, its role in developmental processes such as fruit formation remains poorly understood. Here, we show that the SR splicing factor RS2Z36 is crucial for the early establishment of fruit shape and growth, acting already at the ovary stage and thereby providing a direct link between ovary development and fruit morphology. Our findings demonstrate that RS2Z36 regulates mesocarp cell morphology in the ovary, as two RS2Z36 mutant lines exhibit elongated mesocarp cells, resulting in more elongated ovaries and ellipsoid fruits (Fig. 2-3).

Interestingly, in vegetative organs, RS2Z36 is induced under heat stress and exhibits low expression under control conditions compared to reproductive tissues (Rosenkranz *et al*., 2021). This expression pattern may explain the absence of phenotypes in vegetative organs under control conditions.

A comparative analysis between reproductive and vegetative tissues revealed a higher number of RS2Z36-dependent DAS events and genes in ovaries compared to leaves (Fig. 4; Rosenkranz *et al*., 2024). However, the double *rs2z35 rs2z36* mutant displayed reduced growth rates in vegetative tissues under control conditions (Rosenkranz *et al*., 2024), while the fruits of the double mutant are also significantly smaller (Fig. 2). While RS2Z35 and RS2Z36 are in distinct expression clusters, they might act cooperatively in ovaries (Fig. 1). However, the double mutant did not result in any further change in the fruit index compared to the RS2Z36 single mutant, suggesting that RS2Z36 but not RS2Z35 is responsible for the more ellipsoid phenotype. The contribution of different splicing factors across organs suggests that distinct RNA splicing complexes, assembled from specific SR family members, may operate at different developmental stages, thereby enabling tissue-specific functions. The possibility of differential complex formation is further supported by studies demonstrating specificity in interactions among SR protein pairs, as well as between SR proteins and spliceosome subunits (Golovkin and Reddy, 1998; Tanabe *et al*., 2009; Yan *et al*., 2017). However, detailed investigations on interactions with other SR family members or spliceosome subunits have not yet been conducted.

Similar to leaves, the most affected AS type in ovaries upon *RS2Z36* mutation is A3SS, suggesting a conserved function, although largely on different RNAs (Fig. 4). In addition, similar to leaves, *RS2Z36* mutations caused a misregulation of splicing, meaning that for some introns canonical splicing is enhanced, while for others AS is favoured (Fig. 4). Only in the case of IR, ΔPSI values were shifted predominantly to negative values, suggesting that for this type of splicing event, RS2Z36 functions as intron splicing repressor.

AS has two major effects: it can affect transcript and protein diversity by generating mRNA variants that are translated to protein isoforms, or transcript and protein abundance by shifting RNAs for non-sense mRNA decay (NMD). While some principles for NMD targets have been proposed, two observations suggest that not all transcripts with NMD features are degraded: (i) almost no overlap between DAS genes and DEGs; (ii) identification of peptides that likely derive from transcript variants with long 3’-UTRs, a typical feature of NMD targets. Thus, AS of a subset of genes, e.g. transcription factors and other RNA transcription related genes, can link splicing to changes in gene expression (Fig. 4), as we have previously shown for *HSFA2* (Hu *et al*., 2020). Interestingly, several DEGs code for proteins that are putatively involved in RNA processing, suggesting that there is a level of cross-regulation among the two processes, AS and transcription.

41% of the DAS genes appear to be direct targets of RS2Z36 based on iCLIP data obtained from heat stress leaves in a previous study (Rosenkranz *et al*., 2024). The rest of the genes could be ovary-specific targets or regulated by other splicing factors that are misregulated in the *rs2z36* lines, as for example *SR32* and *SR33* are identified as RS2Z36-dependent DAS in ovaries, while SCL29 was identified as DAP in the proteome analysis. These findings suggest a complex network of interactions where RS2Z36 may influence other splicing regulators, contributing to its global effects on RNA processing.

A manual search in our transcriptome and proteome datasets for genes known to be involved in fruit development (e.g. SUN, OVATE, SlCCS52A, Sl-IAA17, SlGA2ox1, SlLIN5, SlHXK1) did not yield a RS2Z36 target that could confidently explain the phenotypes of the mutant lines. Particularly, we did not find any of the known regulators to be DEG, DAS or DAP, suggesting that RS2Z36 regulates ovary or fruit growth and morphology through different targets. Several candidates could be listed as putative regulators, including DAS genes involved in hormonal signalling (auxin response factors Solyc12006350 and Solyc08g008380; ethylene responsive transcription factor-like Solyc09g059510, AP2-like ethylene responsive transcription factor TOE3 Solyc09g007260), genes related to cytoskeleton (actin-related protein 8 Solyc12g049440), and calmodulin-related genes (Soly10g081170, Solyc02g079040, Solyc12g099340). Further, SlAP2a, a gene previously shown to be involved in fruit ripening and fruit size, but not shape, is an RS2Z36-dependent DAS gene (Chung *et al*., 2010). While histological analysis point to an anisotropic expansion of mesocarp cells and not to cell division as a cause for the elongated fruit phenotype, three cyclin genes were identified as RS2Z36-dependent DAS (Solyc01g044558, Solyc10g078330, Solyc01g107030). Another interesting candidate among DAS genes is Solyc07g008670, annotated as LONGIFOLIA 1 (LNG1). LNG1 regulates leaf morphology by promoting longitudinal polar cell elongation by associating with microtubules (Lee *et al*., 2006). These candidates require further investigation to elucidate their roles in mediating the observed phenotypes in *rs2z36* mutants. Although regulatory events may occur earlier than 2 dba, our choice of this stage is supported by the persistence of the phenotype during fruit development and the higher expression of RS2Z36 at 2 dba compared to anthesis, suggesting that key regulatory genes are already active at this stage.

Cell wall-related modifications are also likely to be related to ovary morphology and consequently fruit size and shape. We identified a pectin acetyltransferase (Solyc10g086480) and a pectin lyase-like protein (Solyc09g075460) as DAS, and a glycine-rich cell wall protein (Solyc12g098225), a hydroxyproline-rich glycoprotein (Solyc03g043740), two chitinases (Solyc04g072000, Solyc10g017980), as well as a pectin methylesterase inhibitor (Solyc06g034370) to be expressed at higher levels in the ovaries of the *rs2z36.1* line compared to WT. Aligning with these findings, we observed an increased signal of LM6-recognized epitopes corresponding to (1,5)-α-L-arabinosyl residues found in the arabinan components of rhamnogalacturonan-I pectic polymers and AGPs in the cell walls of *rs2z36.1* ovaries compared to WT, and a higher abundance of LM20-recognized epitopes indicating elevated levels of partially and highly methylesterified HG (Fig. 6). AGPs are involved in cell expansion (Yang *et al*., 2007) and inhibition of proline hydroxylation, and thereby glycosylation of hydroxyproline-rich glycoproteins including AGPs, results in stunted growth (Fragkostefanakis *et al*., 2014, 2018). Changes in pectin metabolism during the early stages of tomato fruit development have been reported (Fragkostefanakis *et al*., 2012; Terao *et al*., 2013; Kutyrieva-Nowak *et al*., 2024), however, how they contribute exactly to cell differentiation, patterning, division and growth is not understood. Our results point to pectin methylesterification as a central determinant of fruit morphology. Soyama et al. (2025) showed that calcium deficiency reduced LM20 signals and increased LM19 leading to stiffer walls and irregular fruit chambers. In contrast, *rs2z36* ovaries exhibited stronger LM20 and reduced LM19 labelling as well as upregulation of pectin methylesterase inhibitors, suggesting higher levels of methylesterified HG (Fig. 5). This shift could likely alter wall plasticity, resulting in smaller, more ellipsoid fruits. Together, these findings indicate that regulation of cell wall remodelling genes by RS2Z6 may be critical for shaping tomato fruit.

In conclusion, our study establishes RS2Z36 as a key splicing factor controlling fruit growth and morphology in tomato. While we did not identify a straightforward link between gene expression, AS, and protein abundance, our data reveal a highly complex regulatory landscape, underscoring the integrative role of splicing in developmental programs. Importantly, by merging unannotated peptides with putative splice isoforms derived from long-read RNA-seq, we identified novel protein isoforms, highlighting the potential of integrating proteome and transcript isoform data to uncover new layers of gene regulation in plants.

## Materials and Methods

### Plant material

For all experiments in this study, *Solanum lycopersicum* cv. Moneymaker was used as a wild type (WT) or as a background for the generation of mutants. The mutants *rs2z35*, *rs2z36* (here: *rs2z36.1*) and the doublemutant *rs2z35 rs2z36* were described in Rosenkranz *et al*. (2024). The generation of *rs2z36.2* is described below. All plants were grown in the greenhouse with a diurnal cycle of 25°C/16 h with 120 µE illumination and 22°C/8 h without illumination. Plants used for protoplast isolation were grown under sterile conditions in half-strength Murashige and Skoog (MS) medium solidified with GELRITE in Heraeus growth chambers (25°C/16 h with 120 µE illumination and 22°C/8 h without illumination). Mesophyll protoplast and their transformation was performed as described previously (Mishra *et al*., 2002). For immunoblot analysis, 100.000 protoplasts were transformed with a total of 20 µg plasmid DNA (10 µl pRT-3HA-SR; adjusted to 20 µl with a mock plasmid pRT-Neo expressing the neomycin phosphatase coding gene). After PEG-mediated transformation, protoplasts were incubated for 24 h under light (120 µE). To clone pRT-3HA-SR plasmids, the genomic region of *RS2Z36* was amplified by gene specific oligonucleotides (Fw: ACCGGTACCTCCTCGTTATGATGATCGT; Rv: ACCTCTAGAGCGCAAGTTTCAAGGTGACT) from genomic DNA and cloned into a pRT vector together with an 3HA-Tag (HA: human influenza hemagglutinin) via Acc65I/XbaI sites. The generation of the *RS2Z36* WT-CDS encoding plasmid is described in Rosenkranz *et al*. (2021).

### Generation of transgenic plants

For the generation of an additional *rs2z36* line, the same procedure was followed as described in Rosenkranz *et al*. (2024). In brief, two sgRNAs targeting RS2Z36 (gRNA1: GACGGACCCGTTCACGTGATC; gRNA2: GTGATGGACGCCGCATAATTG) were fused to the AtU6 promoter (Addgene #46968) using the MoClo Toolkit (Addgene #1000000044), and combined into the final plasmid pICSL002208 (provided by Dr. Nicola Patron from the Earlham Institute in the UK), which carries the *Streptococcus pyogenes Cas9* gene as well as the kanamycin resistance gene *NPTII*, under the control of the CaMV35S and NOS promoter, respectively. Plant transformation was performed according to McCormick *et al*. (1986) and generated plants were genotyped as described in Rosenkranz *et al*. (2024). Homozygous and T-DNA-free plants were used for analysis.

### Immunoblot analysis

Total proteins were extracted from pelleted protoplasts by adding high salt lysis buffer (20 mM Tris-HCl, pH 7.8; 0.5 M NaCl; 25 mM KCl; 5 mM MgCl_2_; 30 mM EDTA; 0.5% NP40 substitute; 0.2% sarcosyl; 5% sucrose; 5% glycerol; 0.1% ß-mercaptoethanol; supplemented with cOmplete Protease Inhibitor Cocktail, EDTA-free) (Merck) and extracts were treated as described by Port *et al*. (2004). Equal quantities of proteins were separated by sodium dodecylsulfate (SDS)-polyacrylamide gel electrophoresis (PAGE) and transferred onto 0.45 μm nitrocellulose membranes (GE Healthcare). For total protein detection, the membrane was stained with Ponceau Red. Detection of HA-tagged proteins was performed using anti-Hemagglutinin (HA) primary (1:2000, BioLegend) and HRP-conjugated anti-Mouse IgG secondary antibody (1:10,000, Sigma-Aldrich), followed by detection using the ECL Kit (Perkin-Elmer Life Sciences) and the ECL ChemoStar 6 (Intas Science Imaging).

### Morphological and histological analyses

For morphological analysis of the fruit shape and weight, mature fruits were collected and measured using a caliper and a precision scale. For morphological analysis of earlier developmental stages, flowers were cut along the proximal-distal axis, scanned, and the dimensions of fruits and ovaries were measured using the software ImageJ. Floral bud stages were determined based on Shen *et al*. (2019). For cell shape analysis, histological sections of ovaries at 0 days post anthesis (dpa) were prepared according to Pérez-Pérez *et al*. (2023). In brief, ovaries were fixed using 4% paraformaldehyde in phosphate-buffered saline (PBS), dehydrated in acetone, and embedded in Historesin (TECHNOVIT^®^ 8100). Semi-thin sections were cut along the proximal-distal axis using an ultramicrotome (LKB ULTROTOME III^®^), mounted on glass slides and stained with toluidine blue. Pericarp cells were analysed by bright field light microscopy and measured using ImageJ. Fruit- and cell-index was calculated by dividing the maximal length of the fruit or cell by the maximal equatorial diameter.

### Immunofluorescence

Immunofluorescence assays were performed following the procedure described by Pérez-Pérez *et al*. (2019). Semi-thin sections of ovaries (0 dpa: anthesis) were prepared as described for histological analysis, washed with PBS for 1 min, incubated for 5 min in 5% bovine serum albumin (BSA) in PBS, and then treated with the primary rat monoclonal antibody (LM6, LM19 & LM20, PlantProbes, Leeds, United Kingdom) for 1 h. Sections were then washed three times with PBS for 4 min each, incubated 45 min in the dark with secondary antibody (anti-rat IgG, Alexa 488-conjugated, Thermo Fisher Scientific) diluted 1:25 in 1% BSA, and washed three more times with PBS for 4 min each. For DNA staining, DAPI (1 mg/ml) was added to the sections, incubated for 10 min, and washed twice with ddH_2_O for 5 min each. The sections were then mounted with Mowiol and imaged using a confocal microscope (Leica SP5, Leica Microsystems). To allow comparison between the signals, the same excitation and emission-capture settings were applied to each preparation corresponding to the same antibody. Controls were prepared following the same procedure, except the primary antibody was omitted.

### RNA isolation, cDNA synthesis and qRT-PCR

Leaflets or ovaries were homogenized and the RNA was extracted using TRIzol (Invitrogen) following the manufactureŕs instructions. For cDNA synthesis, 1 µg of total RNA was first treated with DNaseI (Roche), following reverse transcription by the RevertAid Reverse Transcriptase (Thermo Fisher Scientific) according to manufactures protocol using oligo(dT)_24_ VN primers. Quantitative real-time PCR (qRT-PCR) was performed using the PowerUp SYBR Green Master Mix (Thermo Fisher Scientific) on a StepOne Plus Real-Time PCR system (Thermo Fisher Scientific) with specific primers for *RS2Z36* (Fw: AATAGCACTCGTCTCTATGTG; Rv: GGATCACTAAATTCTACGAAGG). Relative transcript levels were calculated based on the 2^-ΔΔCt^ method (Livak and Schmittgen, 2001). For normalization, *EF1α* (Fw: TGATCAAGCCTGGTATGGTTGT; Rv: CTGGGTCATCCTTGGAGTT) was used as an internal control. At least 3 biological replicates were performed.

### RNA-Seq and transcriptome analysis

RNA-seq was performed to identify differentially expressed genes (DEGs) in ovaries of WT and *rs2z36.1*. RNA-Seq and downstream transcriptome analyses were performed as described in Rosenkranz *et al*. (2024). Total RNA was extracted in three biological replicates of 10-15 ovaries (2 dba) per sample using TriZol (Invitrogen), followed by DirectZol column purification (Zymo Research) with DnaseI treatment. RNA quality was assessed using the Qubit IQ Assay Kit (Thermo Fisher Scientific) and RNA libraries were constructed from high-quality RNA (IQ score > 8.3) using the NEBNext Ultra II non-directional RNA Library Kit with polyA selection, followed by sequencing as 150 nt paired-end reads on an Illumina NovaSeq 6000 machine, performed by Admera Health, LLC (New Jersey, USA). Initial data processing was carried out using the Galaxy web platform (https://usegalaxy.eu) (Jalili *et al*., 2021). Reads were aligned to the *S. lycopersicum* cv. Heinz 1706 genome (SL4.0 build) and annotated using ITAG4.0 (Fernandez-Pozo *et al*., 2015) with STAR (Galaxy version 2.7.8a) (Dobin *et al*., 2013), allowing soft clipping and up to 5% mismatches. Quantification of gene expression was performed using htseq-count (Galaxy version 0.9.1) (Anders *et al*., 2015). Using the R environment (version 4.1.1) (R Core Team, 2020), removal of low abundance transcripts (sum reads <10 across all samples) was followed by differential expression analysis using DESeq2 (version 1.32.0) (Love *et al*., 2014). Principal component analysis (PCA) was applied to identify batch effects or outlier samples, revealing WT replicate 3 as an outlier, which was excluded from further analysis (Supplemental Figure 4). To control for high dispersion or low counts, log_2_-transformed fold change (log_2_FC) shrinkage was applied with the apeglm package (version 1.14.0) (Zhu *et al*., 2019). Genes were considered significantly differentially expressed if their expression levels in the mutant vs. WT showed |log_2_FC| ≥ 1 and an FDR-adjusted P-value (padj) ≤ 0.05.

### Alternative splicing analysis

Differential AS (DAS) events in ovaries (2 dba) was analysed using MAJIQ and VOILA software packages (version 2.3) (Vaquero-Garcia *et al*., 2016) as described in Rosenkranz *et al*. (2024). First, using *majiq build* on genome-aligned RNA-Seq reads along with the corresponding ITAG4.0 GFF3 annotation file, a splice graph was generated for each gene based on local splicing variations (LSVs). Subsequently, changes in the splice junction usage and intron retention (delta percent selected index, ΔPSI) were quantified and compared between the *rs2z36* lines and the WT using *majiq deltapsi*. *Voila modulize* was employed to reconstruct binary splicing events (e.g., intron retention) from the identified LSVs. This step also calculated the probability of the |ΔPSI| of an LSV surpassing the predefined threshold of 2% (--changing-between-group-dpsi 0.02). In total, 14 distinct binary events were reconstructed, stemming either from single LSVs (e.g. intron retention) or paired LSVs (e.g. cassette exons), whereby *voila modulize* reported two junctions for each LSV. Significance of differential AS events between *rs2z36.1* and WT was determined based on the following criteria: (i) at least one LSV junction pair had |P(|ΔPSI| ≥ 0.02) ≥ 0.9 and |ΔPSI| ≥ 0.05, (ii) all involved junctions had |ΔPSI| ≥ 0.025, (iii) junction pairs of the same LSV exhibited inverse regulation, and (iv) the lower |ΔPSI| value in a junction pair was at least 50% of the higher |ΔPSI| value. Only binary events were analysed (IR, CE, A5SS and A3SS) while complex events were excluded.

### Proteome analysis and identification of novel protein isoforms

High-throughput shotgun proteomics was done according to Chaturvedi *et al*. (2013) with following modifications: Tomato ovaries from WT and *rs2z36.1* were extracted for proteome analysis. Each of 3 biological replicates consisted of ∼17 ovaries (2 dba) which were freeze-dried in liquid N_2_ and ground using mortar and pestle. The proteins were extracted, pre-fractionated (40 µg of total protein were loaded onto a 1D SDS-PAGE gel), trypsin digested and desalted using a C18 spec plate according to previously described procedures (Chaturvedi et al., 2013; Ghatak et al., 2016). One µg of purified peptides was loaded onto a C18 reverse-phase analytical column (Thermo Scientific, EASY-Spray 50 cm, 2 µm particle size). Separation was achieved using a two-and-a-half-hour gradient method, starting with a 4-35% buffer B (v/v) gradient [79.9% ACN, 0.1% formic acid (FA), 20% ultra-high purity water (MilliQ)] over 90 minutes. Buffer A (v/v) consisted of 0.1% FA in high-purity water (MilliQ). The flow rate was set to 300 nL/min. Mass spectra were acquired in positive ion mode using a top-20 data-dependent acquisition (DDA) method. A full MS scan was performed at 70,000 resolution (m/z 200) with a scan range of 380–1800 m/z, followed by an MS/MS scan at 17,500 resolution (m/z 200). For MS/MS fragmentation, higher energy collisional dissociation (HCD) was used with a normalized collision energy (NCE) of 27%. Dynamic exclusion was set to 20 seconds.

Raw data were searched with the SEQUEST algorithm present in Proteome Discoverer version 1.3 (Thermo Scientific) described previously (Chaturvedi et al 2013; Chaturvedi et al., 2015; Ghatak et al., 2020). The identified proteins were quantitated based on total ion count and normalised using the normalised spectral abundance factor (NSAF) strategy (Paoletti et al., 2006). DAPs were identified based on t-test (p ≤ 0.05).

### Splice variant identification via long-read sequencing

RNA libraries were sequenced at Heinrich Heine University (Düsseldorf, Germany). The CCS PacBio Isoseq reads were processed with Iso-Seq tools as follows. After sequencing the barcodes, polyA and concatemers were removed from the ccs reads with lima–isoseq (v1.11.0) and isoseq3 refine (v3.3.0). Afterwards, the reads were clustered and polished with isoseq3 cluster –split-bam 0 and isoseq3 polish. The processed reads were mapped with pbmm2 align –preset ISOSEQ –sort (v1.3.0) against the reference genome *Solanum lycopersicum* cv. Heinz version ITAG4.0. For each sample, sqanti3 generates a gtf file which was analysed by Eventpointer (v 2.4.0, with R v 3.6.3) to get a qualitative overview of splicing events. Eventpointer identifies **AS** events as “Cassette Exon”, “Alternative 3’ Splice Site”, “Alternative Last Exon”, “Retained Intron”, “Alternative 5’ Splice Site”, “Alternative First Exon”, “Mutually Exclusive Exons” and “Complex Event”. Existing databases were used to generate a splice variant atlas (Hosmani et al. 2019). The identified splice variants were merged with the dataset generated in this study using gffcompare (-S –no-merge; v 0.11.6) and used to analyse the common splice variants. Fasta sequences from splice variants were extracted and translated in all three possible forward frames of peptide sequences using the emboss tool transeq -trim -frame F (v. EMBOSS:6.6.0.0).

For each translated frame of each transcript, the ORFs were taken as input for the proteome discoverer to identify potential peptides. The abundant peptides were sorted based on whether they originated from the canonical form or from a splice variant only. To identify proteins with differential abundance between the two treatments, we applied a multiple testing approach. For each ORF, a t-test was performed to assess significant differences, and the resulting p-values were adjusted for multiple comparisons using the Benjamini–Hochberg method.

### Co-expression analysis

Normalized expression values (RPKM) for splicing related genes were obtained by Tomato Expression Atlas (https://tea.solgenomics.net/; Supplemental Table 1). The genes were categorized based on their annotation and orthology analysis as previously shown (Supplemental Table 2; Rosenkranz *et al*., 2021). Clusters for co-expression trends were generated by Mfuzz (Kumar and Futschik, 2007) and visualized as a heatmap with heatmapper (www.heatmapper.ca/).

## Supplementary data

**Supplemental Figure 1.**
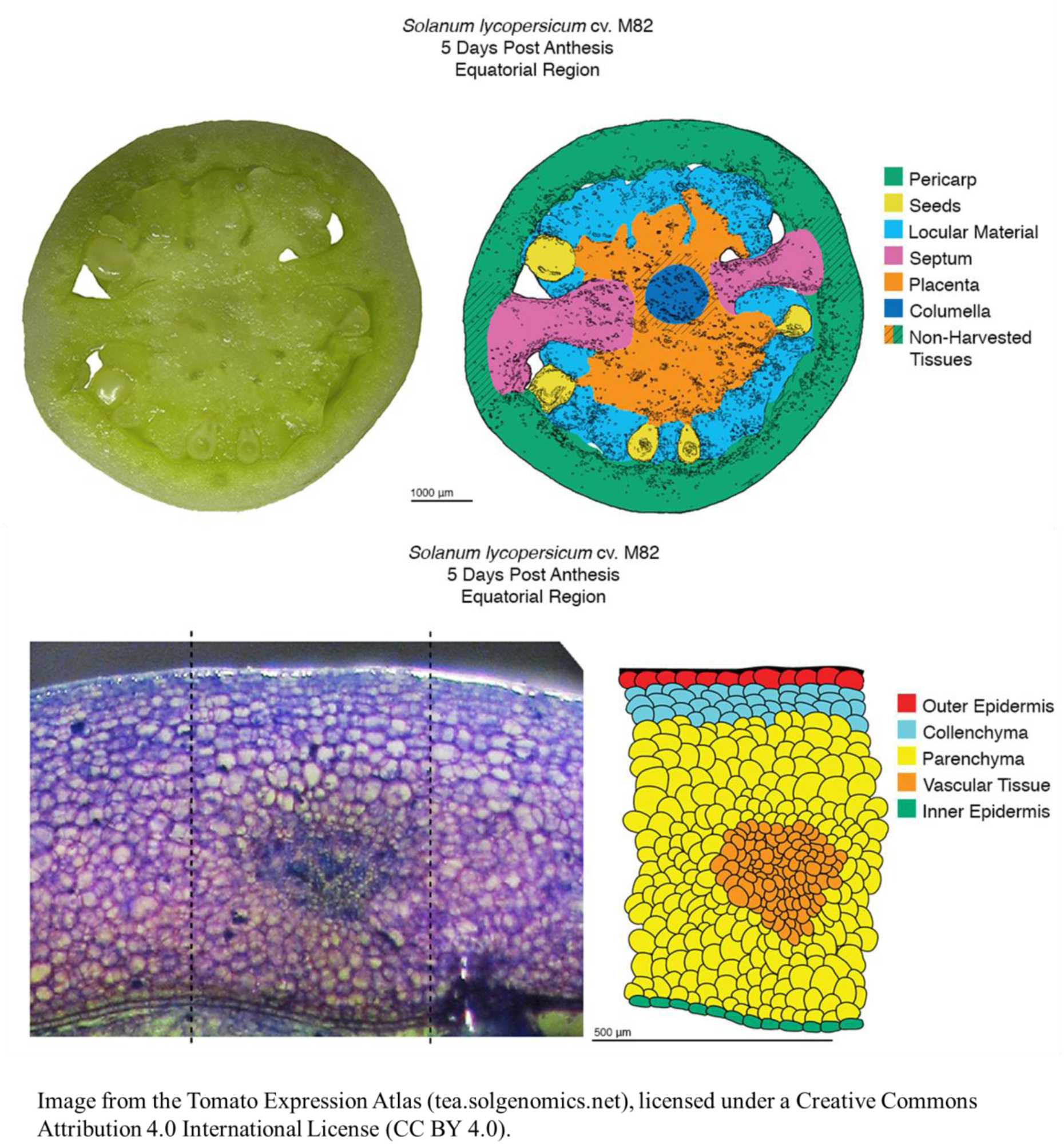
Analysed fruit tissues from TEA database on the example of 5 dpa fruits (Pattison et al., 2015; Fernandez-Pozo et al., 2017; Shinozaki et al., 2018).

**Supplemental Figure 2.**
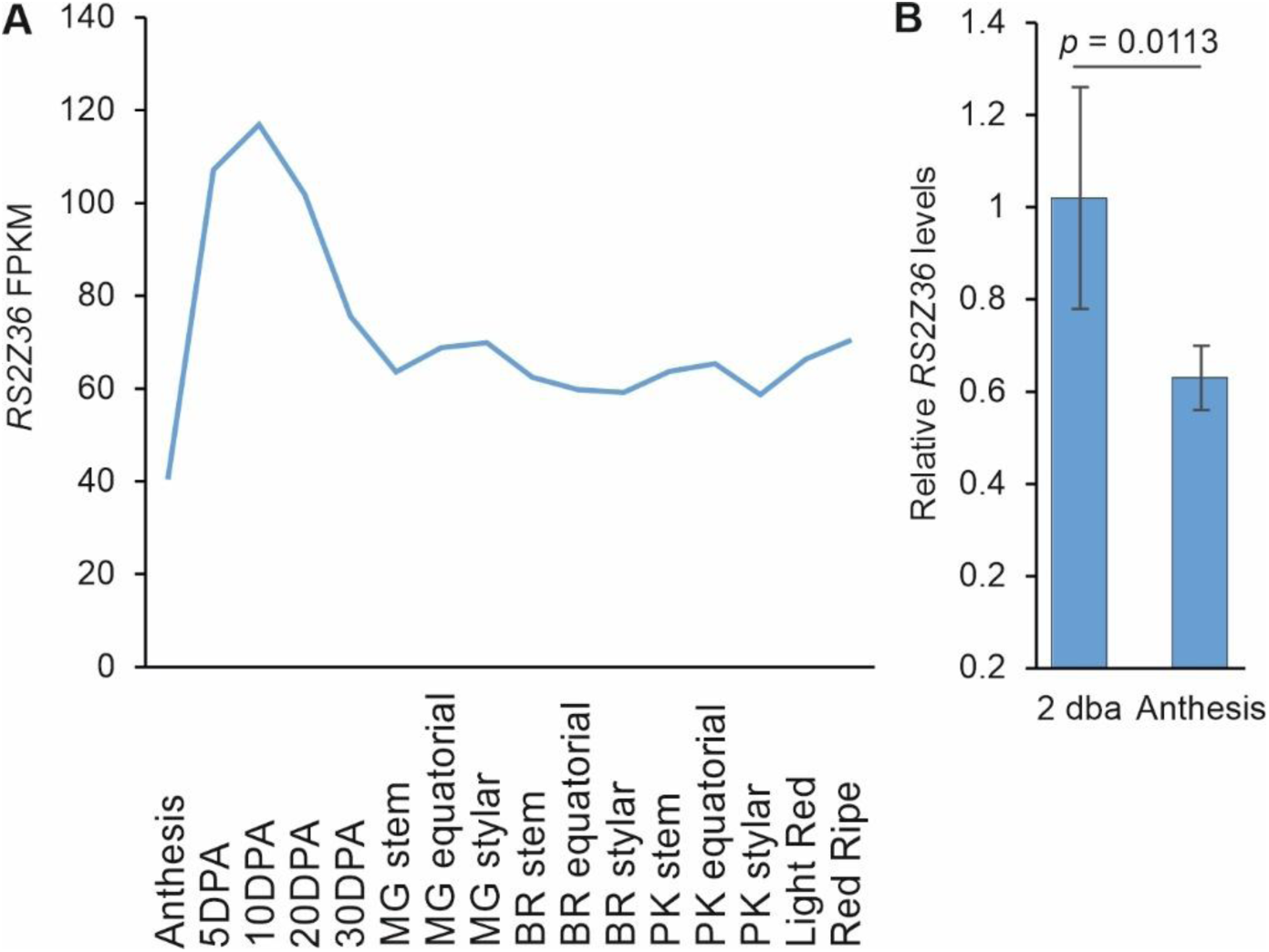
Expression of *RS2Z36* in ovaries and fruits. (A) *RS2Z36* levels based on TEA database (same to Figure 1A). (B) Relative *RS2Z36* levels based on qRT-PCR analysis, in ovaries 2 days before anthesis (dba) and at anthesis. Bars are the average of 5 independent biological replicates ± SD. Statistical significance is based on T-test.

**Supplemental Figure 3.**
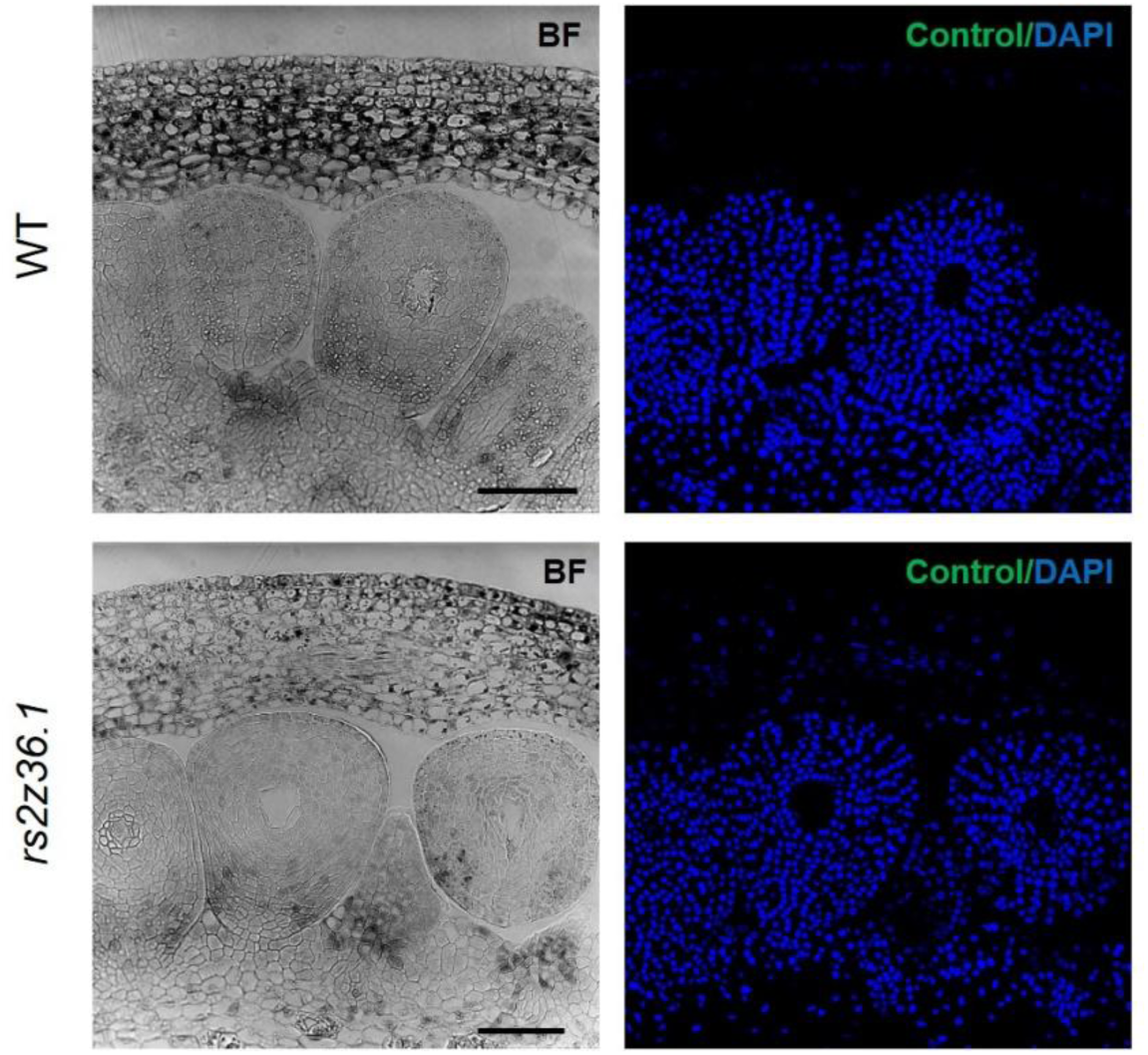
Negative control of the immunohistochemistry samples. Bars are 75 μm.

**Supplemental Figure 4.**
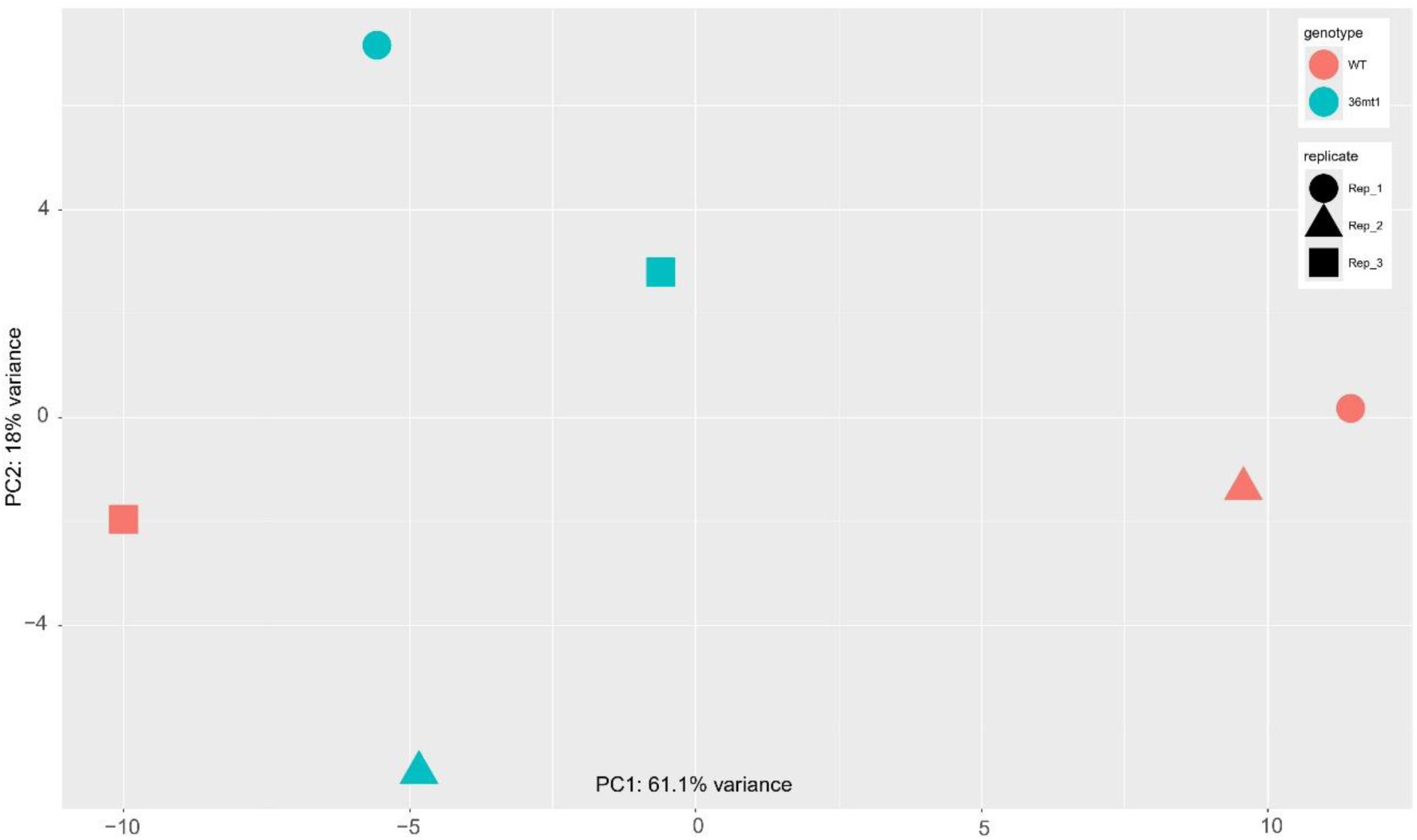
PCA of RNA-seq samples based on top 500 most variable genes.

**Supplemental Figure 5.**
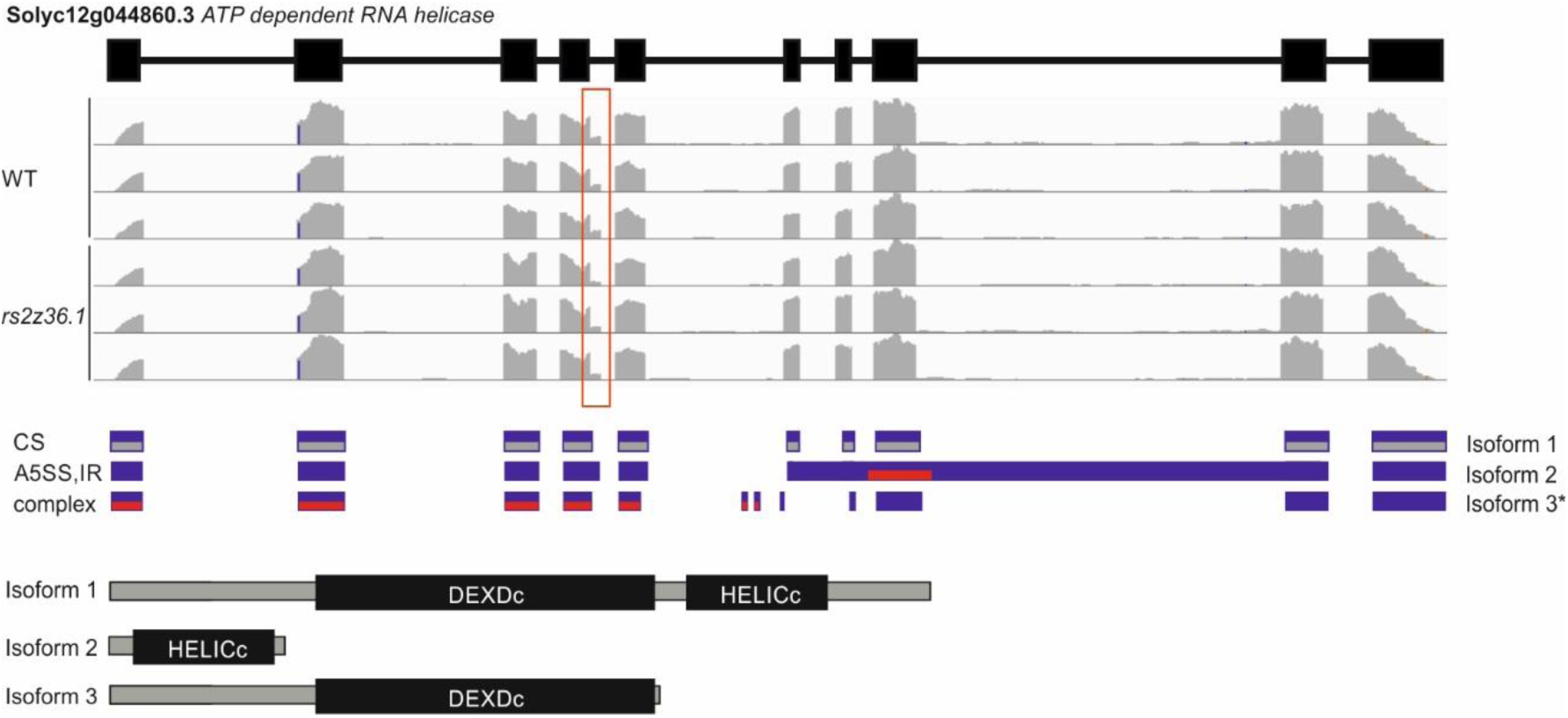
Gene structure, read coverage, RNA splice variants, and protein isoforms with the indicated domains of *Solyc12g044860* gene. Grey boxes in the splice variants indicate the coding regions. Red boxes indicate the coding regions for the identified unannotated peptides. Violet colour indicates RNA variants. The asterisk indicates a splice variant identified in the Iso-seq library but is not an RS2Z36-dependent DAS event.

Supplemental Table 1. Expression (RPKM) of splicing related genes in different tomato fruit tissues across different developmental stages.

Supplemental Table 2. Categories of splicing related genes.

Supplemental Table 3. List of all identified differentially splice events in wild type and *rs2z36.1* ovaries. Supplemental Table 4. List of differentially expressed genes in wild type and *rs2z36.1* ovaries.

Supplemental Table 5. Mapman categories of DAS genes and DEGs.

Supplemental Table 6. Differentially abundant proteins (annotated) in wild type and *rs2z36.1* mutant ovaries.

Supplemental Table 7. Identified peptides corresponding to novel protein isoforms derived from RNA splice variants in wild type and *rs2z36.1* mutant ovaries.

Supplemental Table 8. MapMan categories of annotated DAP and DAP isoforms in wild type and *rs2z36.1* mutant ovaries.

## Acknowledgements

We thank Holger Schranz for the cultivation of plants, Daniela Bublak for the technical support and Enrico Schleiff for the valuable discussions. SV acknowledges the support with a Short Term Scientific Mission by the COST Action EPICATCH (CA19125). SF, SS and PC acknowledge network opportunities by RECROP COST Action (CA22157). This work has been supported with a grant by DFG (FR 3776/5-1) to SF.

## Author contributions

SF, SV, RRER: conceptualization; SV, RRER, MK, YPP, SB, KJZ, JB, SS, PC, AG, LAS: methodology; SF, RRER, MK, JB, SS, PC, KZ, AG, LAS: formal analysis; SV, RRER, MK, YPP, SB, KZ, JB, SS, PC: investigation; SF, SS, PC, PST, KJZ, WW, LAS: resources; SF, SV, RRER, MK, SS, PC, KJZ: data curation; SF, SV, RRER: writing - original draft; SV, RRER, MK, YPP, SB, KZ, JB, SS, PC, PST, MMM, KZ, SF: writing - review & editing; SV, RRER, YPP, SF: visualization; SF, SS, PST, MMM, KZ: supervision; SF, PC: funding acquisition

## Data availability

The RNA-Seq data underlying this article are available in the NCBI Gene Expression Omnibus (GEO) Database at www.ncbi.nlm.nih.gov/geo/ with accession number GSE286889. Proteome data is available in PRIDE database (https://www.ebi.ac.uk/pride/) with the accession number PXD060312.

